# Adenosine A1R/A3R Agonist AST-004 Reduces Brain Infarction in Mouse and Rat Models of Acute Ischemic Stroke

**DOI:** 10.1101/2022.03.14.484307

**Authors:** Elizabeth S. Fisher, Yanan Chen, Mikaela M. Sifuentes, Jeremy J. Stubblefield, Damian Lozano, Deborah M. Holstein, JingMei Ren, Nicholas DeRosa, Tsung-pei Chen, Gerard Nickel, Theodore E. Liston, James D. Lechleiter

## Abstract

Acute ischemic stroke (AIS) is the second leading cause of death globally. No Food and Drug Administration (FDA) approved therapies exist targeting cerebroprotection following stroke. Our group recently reported significant cerebroprotection with the adenosine A1/A3 receptor agonist, AST-004, in a transient stroke model in non-human primates (NHP) and in a preclinical mouse model of traumatic brain injury (TBI). However, the specific receptor pathway activated was only inferred based on *in vitro* binding studies. The current study investigated the underlying mechanism of AST-004 cerebroprotection in two independent models of AIS: permanent photothrombotic stroke in mice and transient middle cerebral artery occlusion (MCAO) in rats. AST-004 treatments across a range of doses were cerebroprotective and efficacy could be blocked by A3R antagonism, indicating a mechanism of action that does not require A1R agonism. The high affinity A3R agonist MRS5698 was also cerebroprotective following stroke, but not the A3R agonist Cl-IB-MECA under our experimental conditions. AST-004 efficacy was blocked by the astrocyte specific mitochondrial toxin fluoroacetate, confirming an underlying mechanism of cerebroprotection dependent on astrocyte mitochondrial metabolism. An increase in A3R mRNA levels following stroke suggested an intrinsic cerebroprotective response that was mediated by A3R signaling. Together, these studies confirm certain A3R agonists, such as AST-004, are promising new therapeutics for the treatment of AIS.

## Introduction

Acute ischemic stroke (AIS) occurs in ∼795,000 individuals annually in the US, resulting in over 147,000 deaths, and often permanent disability in those who survive^1^. Globally this number rises to over 6.5 million annual deaths^2^. Current approved treatments are limited and focus only on restoration of cerebral blood flow to the ischemic area of the brain, achieved by either intravenous administration of tissue plasminogen factor (tPA) to dissolve blood clots and/or endovascular mechanical thrombectomy to remove a large vessel blood clot causing ischemia^3^. These therapies are constrained to small subsets of patients in the emergency departments of both primary and comprehensive stroke centers; ranging from 10 - 14% of patients for tPA and from 1 - 4 % of patients for endovascular clot removal, respectively^4^. Moreover, neither therapy directly targets neuronal survival within the ischemic penumbra, which if rescued would lead to better patient outcomes^5,6^.

Loss of oxygen and glucose during ischemia dysregulates many energy-dependent processes in the brain, leaving affected tissue at significant risk of damage and cell death^7^. While brain tissue at the core of an infarction is rapidly and irreversibly lost, cells in the surrounding hypoperfused penumbra die over hours and days after the initial ischemic event, and thus have the potential to be rescued^8,9^. Research by our group demonstrated the intrinsic healing mechanisms of astrocytes could be enhanced by improving their mitochondrial ATP production, leading to an energy-dependent reduction in the size of brain lesions in mouse models of stroke^10,11^. This early research focused on P2Y_1_ receptor activation (P2Y_1_R), using the P2Y_1/12/13_ receptor agonist 2-methylthio-adensoine di-phosphate (2-MeSADP) and then the specific P2Y_1_R agonist MRS2365. Subsequent research on the pharmacokinetics (PK) and metabolism of these compounds indicated MRS2365 was a prodrug that was rapidly and completely metabolized *in vivo*, to the novel nucleoside metabolite AST-004^12^. Cerebroprotection with these first-generation purinergic agonists^10,11,13^ was attributed to the conformationally constrained metabolite of MRS2365, AST-004, and the non-conformationally constrained metabolite for 2MeS-ADP, 2-methylthioadenosine^12^.

We recently confirmed the cerebroprotective efficacy of AST-004 treatments in a preclinical mouse model of traumatic brain injury (TBI) and a transient stroke model in non-human primates (NHP)^14^. However, the specific receptor pathway activated was only inferred from binding studies showing AST-004 interacted with the adenosine A3 receptor (A3R), with some affinity for the adenosine A1 receptor (A1R)^12^. Here, we further investigated the underlying mechanism of AST-004 cerebroprotection in two independent models of AIS: permanent photothrombosic stroke in mice and transient middle cerebral artery occlusion (MCAO) in rats. We found AST-004 treatments were effective across a range of doses and that cerebroprotection could be blocked by the A3R antagonist MRS1523. The high affinity A3R agonist MRS5698 also decreased infarct size in the mouse photothrombotic model, whereas the high affinity A3R agonist Cl-IB-MECA was ineffective under our mouse photothrombosis experimental conditions. Cerebroprotection was blocked by the astrocyte specific mitochondrial toxin fluoroacetate, indicating a mechanism of action dependent on astrocyte mitochondrial ATP production. We also found increased levels of A3R mRNA following stroke, revealing an intrinsic protective response that could be exploited for treatment. A3R agonists have been reported as cerebroprotective against stroke since the mid-1990s^15-25^, and most recently for TBI^26,27^. Together, these studies confirm that certain A3R agonists, such as AST-004, may be exciting new treatment options that need to be developed for clinical evaluation.

## Experimental Procedures

### Animals

Male and female C57BL/6 mice with access to food and water ad libitum were housed in 12-hour light-dark cycles. Experimenter was blinded to experimental groups. All mouse experiments were performed in accordance with the Institutional Animal Care and Use Committee at UT Health San Antonio. All rat surgeries were performed at NeuroVasc Preclinical Services Inc. (Lexington, MA). Preclinical services and procedures reviewed and approved by the IACUC at NeoSome (Lexington, MA). Male Wistar rats 225-250g were ordered 7-10 days prior to surgery (Charles River Laboratories, Wilmington MA). They were allowed free access to food and water and housed two per cage.

### Photothrombotic stroke

Photothrombic stroke was induced using the protocols as previously described^10,11^. Briefly, mice were anesthetized with 4% isoflurane and maintained at 2% isoflurane throughout the surgery. Hair was removed, and incision made on the dorsal scalp, and head mounted in a custom frame. Either a cranial window or a thin skull prep was performed. Rose Bengal (Sigma, Cat no 330000) dye was then injected intravenously, and a blood clot induced with a 568 nm laser on a Nikon (TE 200) microscope, with blood vessels between 30-40 μM targeted for clotting. Mice were injected with drugs (MRS2365: Tocris Cat no 2157; MRS1523: Sigma Cat no M1809; Fluoroacetate: Sigma Cat no 62-74-8; MRS5698: Tocris Cat no 5428; Cl-IB-MECA: Tocris Cat no 1104; AST-004, synthesized at the National Institute of Diabetes, Digestive and Kidney Diseases, Bethesda, MD^28^) either before surgery or 30 minutes post-stroke, as described in the paper.

### TTC Staining and lesion volume quantification

2,3,5-Triphenyltetrazolium chloride (TTC, Sigma Aldrich, Cat no T8877) staining was performed as previously described. Briefly, brains were removed and placed in ice-cold PBS for 5 minutes on ice. Then brains were placed in a tissue matrix (Ted Pella, Cat no 15050), and sliced 1-mm thick coronal section through the brain and placed in solution with TTC for 16 minutes at 37 degrees, turning over halfway through incubation. Once stained, sections were fixed overnight with 4% PFA.

Stained and fixed sections were placed on a scanner with a ruler and images exported to ImageJ for analysis. Area was quantified by measuring white sections using the freehand tool of ImageJ of tissue visible on the slice, and volumes from each section were added to make the total lesion volume.

### Western blotting

A 1 mm punch was removed each from ipsilateral and contralateral sides of a stroke brain. The punch was then sheered in sample buffer and sonicated for 10 minutes and spun at 10,000 rpm. Supernatants were collected and 50 μg of protein was loaded onto a 10% SDS gel and run and transferred to a nitrocellulose membrane. Blots were then probed for GFAP (Dako Omnis Cat no Z0334) at 1:1000 and GAPDH (Cell Signaling Cat no 97166S) 1: 1000. LiCor secondary antibodies (donkey anti-rabbit Cat no 926032213 and donkey anti-mouse Cat no 926-68022) were used and images were developed using the Odyssey system.

### RNA isolation and Quantitative real time-PCR (qRT-PCR)

#### Tissue Harvest

24 hours after stroke, mice were euthanized, and their brains were rapidly removed, and 1 mm serial coronal sections were collected. Following TTC staining and scanning, brain sections were placed under a dissection microscope and the region of tissue classified as “stroke lesion” (i.e. the section of the brain that remained white following incubation in the TTC solution) was removed through cutting with a scalpel or pair of fine dissection scissors (Fine Science, Cat no 15018-10). The stroke lesion samples were pooled in a 1.5 mL microcentrifuge tube, weighed, and then flash-frozen in liquid nitrogen. The side of the brain with a stroke lesion was termed “ipsilateral.” An equivalently sized piece of tissue was cut from the opposite side of the brain (contralateral) on each section that displayed a stroke lesion. The contralateral pieces were also pooled in a separate 1.5 mL microcentrifuge tube, weighed and flash-frozen in liquid nitrogen. Tubes were then stored in a -80°C freezer for down-stream processing.

#### RNA Isolation and cDNA synthesis

Frozen ipsilateral and contralateral tissue was ground into a tissue powder using a liquid nitrogen mortar and pestle. The tissue powder was transferred to 2 mL microcentrifuge tube and 1 mL TRIzol Reagent (ThermoFisher, Cat no 15596018) was added. The tissue powder sample was homogenized for 15-20 seconds in TRIzol using a mechanical hand-held homogenizer. Following homogenization, samples were centrifuged at 12,000 x g for 10 minutes in a 4° centrifuge and the supernatant was transferred to a separate tube for RNA isolation. The RNA isolation procedure was then followed according to the manufacturer’s instructions. Isolated RNA was resuspended in DEPC-treated ddH_2_O and checked for concentration and quality of a Nanodrop 2000. RNA samples were diluted to 200 ng/μL and 1 μg of RNA was converted into cDNA using a High-Capacity cDNA Reverse Transcription Kit according to the manufacturer’s instructions (ThermoFisher, Cat no 4368814). cDNA was then diluted 1:5 in ddH_2_O for downstream analysis.

#### qRT-PCR

Diluted cDNA (1:5 dilution) was used for quantitative Real Time-PCR (qRT-PCR). Genes of interest were normalized to *Gapdh* gene expression. Expression was determined using the ΔΔCT method. Primer sequences were designed using Primer-BLAST (http://www.ncbi.nlm.nih.gov/tools/primer-blast/). Primer sequences were as follows: *Gapdh* (Forward 5’-3’ CAAGGAGTAAGAAACCCTGGACC, Reverse 5’-3’ CGAGTTGGGATAGGGCCTCT) ^29^, *Gfap* (Forward 5’-3’ AAAACCGCATCACCATTCCTG, Reverse 5’-3’ GTGACTTTTTGGCCTTCCCC) and *Adora3* (Forward 5’-3’ GACAGTCAGATATAGAACGGTTACCAC, Reverse 5’-3’ TTCCAGCCAAACATGGGGGTCA). qRT-PCR amplification of target genes was achieved using Power SYBR Green PCR Master Mix (ThermoFisher, Cat no 4368706) and primer forward/reverse mixes at a final primer concentration of 150 nM in a 10 μL reaction on a 384-well plate. qRT-PCR was conducted on an Applied Biosystem 7900HT Real-Time PCR System.

### Rat tMCAO Surgeries

Briefly, the middle of the neck was shaved with electric clippers and cleaned with Hibiclens. A skin incision was made over the right common carotid artery (CCA), the muscle was retracted, and the CCA bifurcation was exposed. The CCA was ligated and, and a distal segment of the external carotid artery (ECA) was temporarily clamped using a suture or clip. A nylon suture was then inserted through the CCA and advanced into the internal carotid artery (ICA) for a predetermined distance (18-21 mm) based on animal weight. The ECA clip/suture was then removed. After a cannulation of the right jugular vein with PE-90 tubing, the skin incision was closed with surgical staples. The animal was again anesthetized after the 90 minutes ischemic period, the wound was re-opened, and the intravascular suture was removed from the CCA, initiating reperfusion. The bolus dose, followed by the primed pump connection, was administered immediately upon suture removal and the skin wound was again closed. During the time of anesthesia, a self-regulating heating pad connected to a rectal thermometer was used and maintained at 37.0° ± 1°C. Cefazolin (40 mg/kg; Hikma Parma Corp 156005) was given intraperitoneally. before surgery to prevent infections. Subcutaneous buprenorphine (∼0.1 mg/kg; Par Pharm companies, Inc.) was given before surgery as analgesia. All treatment solutions were stored in 4°C and kept in 4°C until use. Vehicle: DMSO, High Dose Group 3 mg/kg bolus and 0.042 mg/min/kg Alzet Infusion, Mid Dose Group 0.4 mg/kg bolus and 0.0056 mg/min/kg Alzet Infusion, Low Dose Group 0.04 mg/kg bolus and 0.00056 mg/min/kg Alzet Infusion. Starting at the time of reperfusion, animals received 1 ml/kg intravenous bolus followed by Alzet Infusion (8 μl/hr for 24 hours) through jugular vein. These dosing regimens were designed to maintain targeted steady-state plasma and brain concentrations of AST-004 throughout the evaluation period. Treatments were randomly assigned to each day of surgery: (https://www.randomizer.org).

### Rat functional assay

Functional activities were evaluated using modified neurological rating scale (mNRS)^30^. Modified Neurological Rating Scale (mNRS): 0 - Indicated no neurologic deficit; 1 - Failure to extend left forepaw fully, a mild focal neurologic deficit; 2 - Circling to the left, a moderated focal neurologic deficit; 3 - Falling to the left, a severe focal deficit: 4 - Rats did not walk spontaneously and had a depressed level of consciousness. 5 – Death.

### Rat sacrifice and lesion volume quantification

Twenty-four hours after reperfusion/dosing, rats were sacrificed using CO_2_, and brains were removed and cut into seven 2 mm thick coronal sections using a rat brain matrix (+4.7, +2.7, +0.7, -1.3, -3.3, -5.3 and -7.3, compared to bregma respectively) and stained with TTC. The brain sections were put into 2% TTC solution in a dark place at room temperature for 30 minutes. The TTC solution was then changed to 10% formalin for fixation until images were captured approximately 24 hours later. Images were captured using a digital camera fixed on a photo stand. Volumetric analysis of the infarct area was performed using Image J (NIH software). The free-hand tool was used to trace the area of the infarcted tissue of the right hemisphere, the uninfarcted tissue for both hemispheres. Infarct area was calculated by subtracting the uninfarcted area of the ipsilateral hemisphere from the area of the intact contralateral hemisphere. The volume of each hemisphere was then calculated by multiplying the area with section thickness (2 mm) and the number of sections in between each sampling (7). The infarct volume was expressed as a percentage of the intact hemispheric volume.

### Statistics

Statistical analysis was performed in GraphPad/Prism. All data were expressed as mean ± S.E.M. All pairwise comparisons were performed using student’s t-test. One way-ANOVA was used in the multiple dosing experiments, and for examining inter-sex differences. Rat data was analyzed by one way ANOVA.

## Results

### AST-004 is cerebroprotective in mice after permanent photothrombotic occlusion over a wide range of doses

To test whether the A1R/A3R agonist, AST-004, was protective against cerebral ischemia, we induced stroke using photothrombosis and intraperitoneally (IP) injected AST-004 within 30 minutes of stroke onset. Twenty-four (24) hours post-stroke, brains were harvested, coronally sliced into 1 mm sections, then stained with triphenyltetrazolium chloride (TTC), a dye which turns red under dehydrogenase activity in live tissue. Cortical tissue near the site of injury that remained white was considered part of the ischemic lesion. Lesion volumes were estimated by serially integrating the lesions areas in each 1 mm-thick brain section. Control mice injected with vehicle alone exhibited an average lesion volume of 12.59 +/-1.56 mm^3^ (n = 23) (**Fig. 1 A**). We tested 3 AST-004 doses a full log difference from each other: low (0.022 mg/kg), mid (0.22 mg/kg), and high (2.2 mg/kg) concentrations. We found that all 3 doses significantly decreased lesion volume (**Fig. 1 B, C**). The low dose of AST-004 (L) reduced lesion size to 9.26 +/-1.67 mm^3^ or 63.38 +/-10.87% of vehicle treated mice (n = 15, p < 0.05). Those mice treated with the middle AST-004 (M) dose following photothrombotic stroke showed significantly reduced lesion volumes with an average of 5.92 +/-0.88 mm^3^ or 48.39 +/-6.59% of vehicle (n = 24, p < 0.001), while the high dose of AST-004 (H) reduced the average lesion size to 8.80 +/-1.48 mm^3^ or 60.38 +/-8.92% of vehicle (n = 14, p < 0.04) (**Fig. 1 B, C**). The composite lesion size when all 3 doses of AST-004 were pooled was 7.63 +/-0.75 mm^3^ or 66.05 +/-5.67% of vehicle (n = 53, p<0.006).

**Figure 1:**
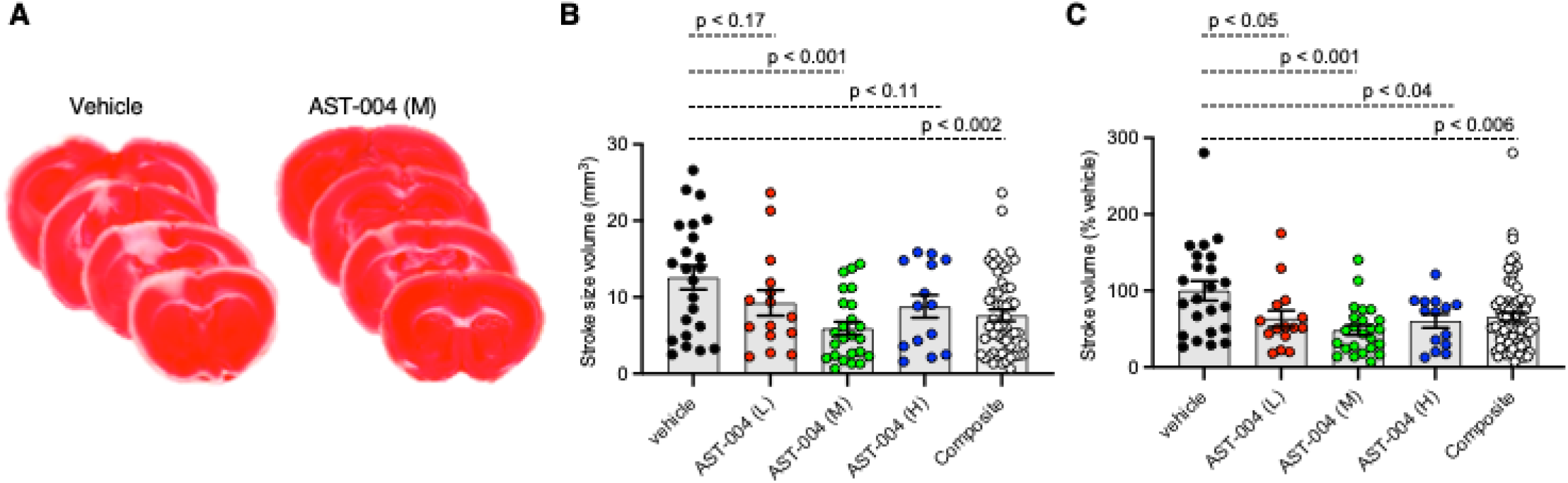
Photothrombosis-induced stroke infarctions are reduced by AST-004 treatments. **(A)** Coronal sections of brains stroked with photothrombosis for vehicle (saline injected) and AST-004 (mid dose) treated mice. **(B)** Average TTC-stained stroke volumes in mice treated with either vehicle or AST-004 at the concentrations labeled: 0.022mg/kg (L), 0.22mg/kg (M) and 2.2mg/kg (H). Composite group are all values from L, M and H AST-004 doses. **(C)** Data in panel B normalized to the mean vehicle lesion size and replotted. Data were pooled from 3 experiments using male mice and plotted as mean +/-SEM.

### AST-004 is cerebroprotective in mice following stroke in females and males

To assess whether AST-004 efficacy was sex-dependent, we re-plotted the data presented in Figure 1, separating lesion data into groups for male and female mice. We found no significant differences between female and male average lesion sizes, 9.48 +/-2.40 mm^3^ (n = 10) vs 15.04 +/-1.89 mm^3^ (n = 13), respectively in vehicle treated mice. Lesion sizes for AST-004 treated mice were significantly reduced for females, 3.95 +/-0.95 mm^3^ (p < 0.029, n = 13) or 37 +/-8.18 % of vehicle and also for males, 8.25 +/-1.25 mm^3^ (p < 0.009, n = 11) or 56.20+/-7.91 % vehicle, indicating sex was not a determinant for AST-004 efficacy (**Fig. 2A, B**). We note that although male lesion sizes trended higher, even when expressed as a % of the mean vehicle lesion size, these differences were not statistically different (**Fig. 2A, B**).

**Figure 2:**
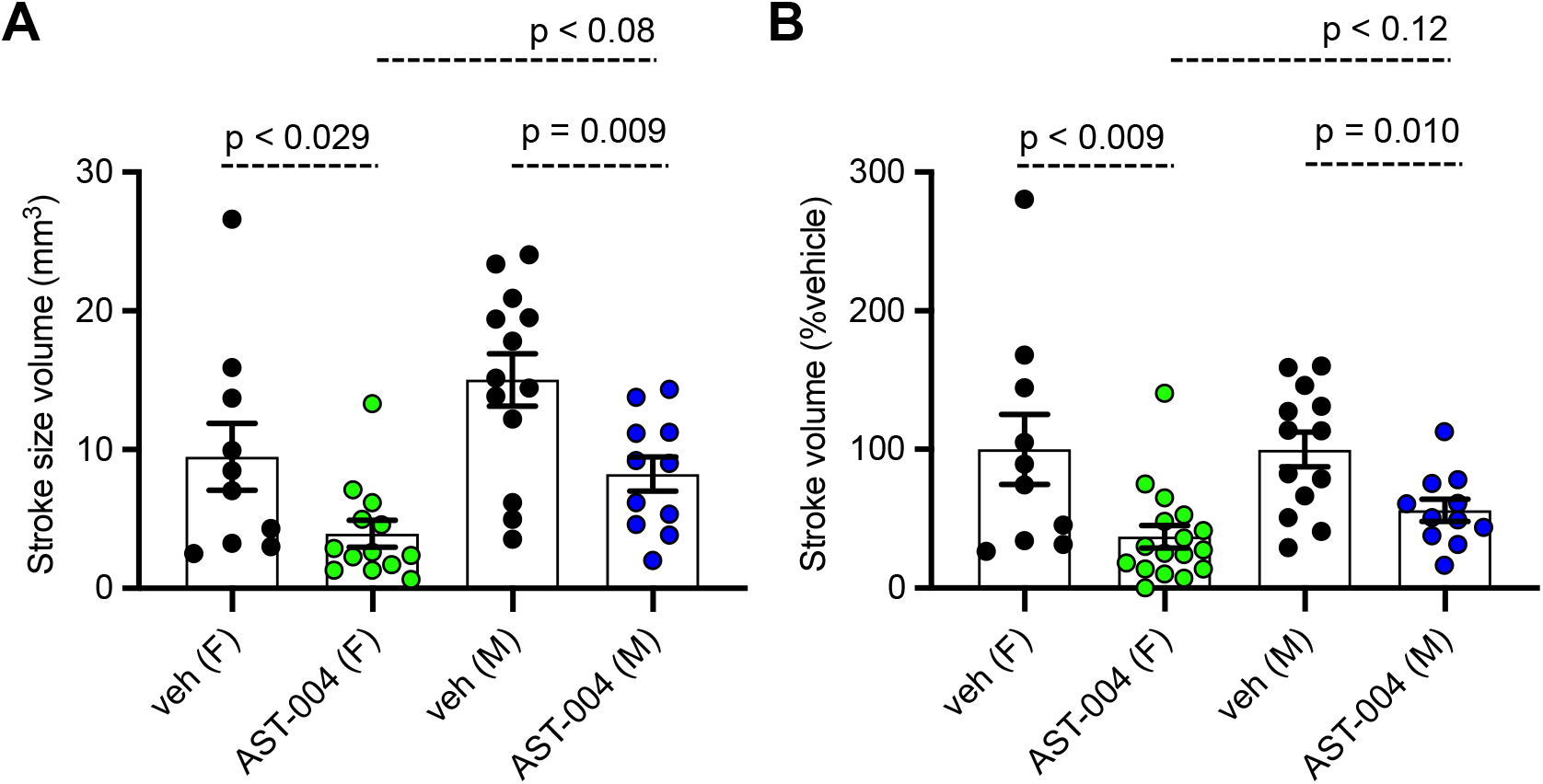
Photothrombosis-induced stroke infarctions are reduced by AST-004 in both male and female mice. **(A)** Average TTC-stained stroke volumes for male (M) and female (F) mice. **(B)** Female and male groups replotted as a % of their vehicle-treated means. Data were pooled from 3 experiments and plotted as mean +/-SEM.

### AST-004 cerebroprotection is blocked by an A3R antagonist

We recently reported AST-004 was primarily a moderate affinity A3R (human K_i_ 1490 nM) and A1R (human K_i_ 1590 nM) agonist^12^. For comparison, the natural endogenous ligand, adenosine, has Ki values of 70 nM and 6500 nM for human A1Rs and A3Rs, respectively^31^. This equates to ∼4x higher affinity of AST-004 for human A3Rs and to ∼23x lower affinity of AST-004 for human A1Rs compared to adenosine. Accordingly, we tested whether cerebroprotection was inhibited by pre-injecting mice with the A3R antagonist, MRS1523 (2 mg/kg), 15 minutes before inducing a photothrombotic stroke. MRS1523 (2 mg/kg) was also added along with vehicle or AST-004 after each stroke. Brains were harvested 24-hours post stroke, sliced, stained for TTC, and lesion volumes measured as described in Figure 1. We found that MRS1523 by itself, did not significantly affect the average lesion volume (**Fig. 3 A, B**). The mean lesion size was 11.57 +/-2.90 mm^3^ (n = 7), comparable to the average lesion size for mice treated with only vehicle (dashed line, **Fig. 3 B**). In contrast, MRS1523 completely blocked AST-004 efficacy. The average lesion size was 14.48 +/-1.60 mm^3^ (n = 8, pooled from low and high AST-004 treated mice), not significantly different than the average lesion size in control mice. For comparison, the average lesion size in mice treated with AST-004 alone is shown as the dashed red line (**Fig. 3 B**). We conclude from these data that AST-004 cerebroprotective efficacy requires activation of the A3R.

**Figure 3:**
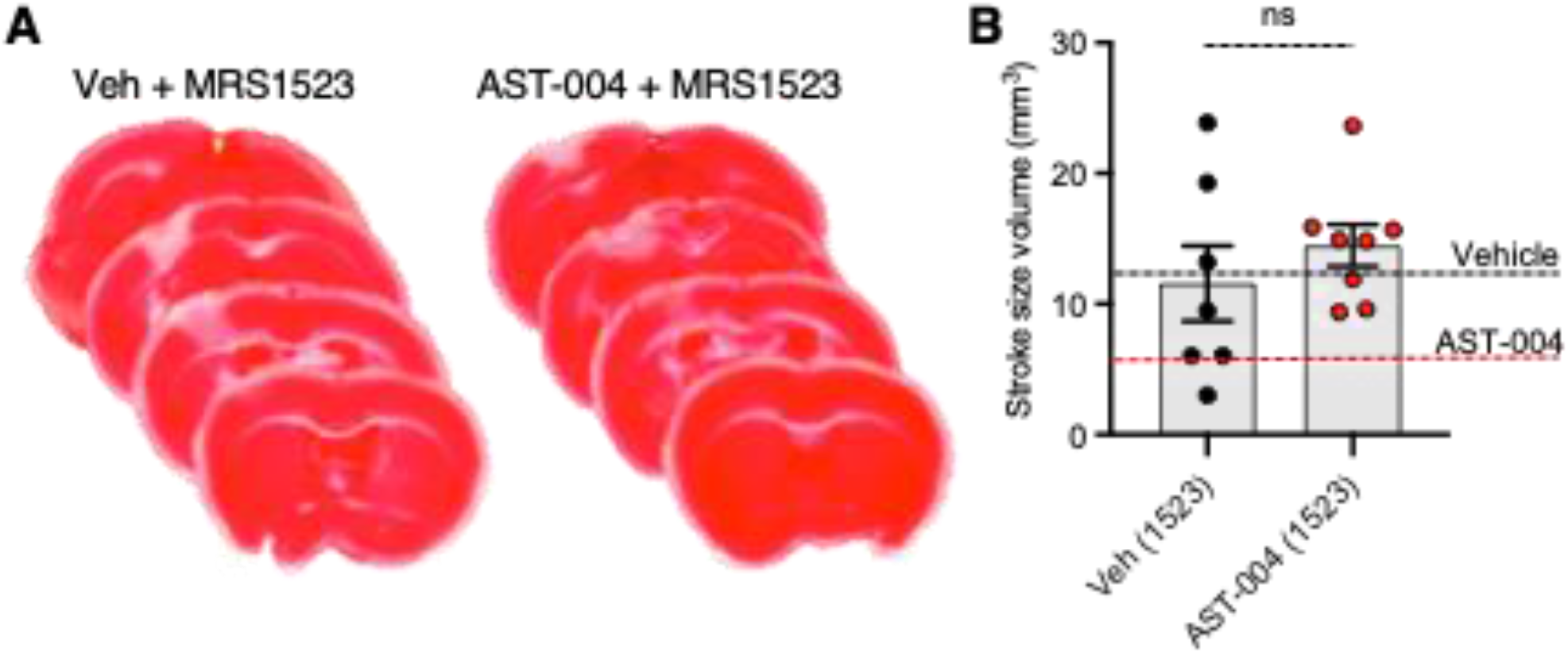
Photothrombosis-induced stroke infarctions reduced by AST-004 are blocked by A3R antagonist MRS1523. (A) Coronal sections from stroked mice pre-injected 15 minutes before stroke onset with the A3R antagonist MRS1523, stroked and post-treated with vehicle or AST-004. (B) Average TTC-stained stroke volumes in mice treated with MRS1523 alone or with AST-004 and MRS1523.

### A3R agonist MRS5698, but not Cl-IB-MECA is cerebroprotective

We also tested the efficacy of the high affinity A3R agonists MRS5698 and Cl-IB-MECA. MRS5698 treatments (1.4 mg/kg) significantly reduced the mean lesion volume (7.33 +/-1.07 mm^3^, n=11) compared to untreated mice, although this reduction lesion size appeared smaller than that observed for AST-004 (**Fig. 4**). However, in mice treated with the A3R agonist Cl-IB-MECA (0.19 mg/kg), the mean lesion size 24 hours post-stroke (16.06 +/-3.18, n = 5) trended higher than untreated mice (**Fig. 4**). This may be due to the low blood barrier permeability of this compound or low unbound brain fraction available to interact with A3R. These data suggest that A3R agonism by itself, may not be sufficient for cerebroprotection as both MRS5698 and Cl-IB-MECA have substantially higher affinity for A3R than does AST-004, but this higher affinity does not translate to higher efficacy.

**Figure 4:**
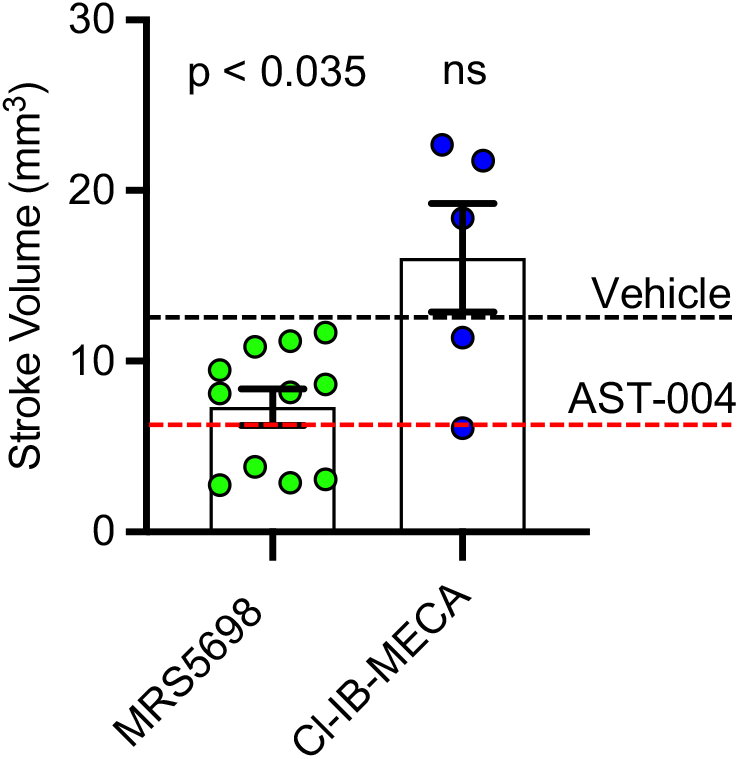
Cerebroprotective efficacy of A3R agonists MRS5698 and Cl-IB-MECA in photothrombosis-induced stroke infarctions. Average TTC-stained stroke volumes as labeled. Black dashed line shows mean stroke volume for untreated mice. Red dashed line shows mean stroke volume for AST-004 treated mice as presented in Fig. 1B. Statistics relative to vehicle control in Fig. 1B. Data were pooled from 2 experiments (N = number of mice per treatment) and plotted as mean +/-SEM

### AST-004 exerts its cerebroprotective effects through astrocyte energy metabolism

Earlier studies by our group indicated MRS2365 cerebroprotection was the result of enhanced mitochondrial energy metabolism in astrocytes^10,11^. To test whether the cerebroprotection efficacy of AST-004, a MRS2365 metabolite, was similarly dependent on mitochondrial metabolism, we pre-treated mice with the astrocyte specific mitochondrial toxin, fluoroacetate (FAc, 0.004mg/kg)^32^. Consistent with the date presented above, treatment of mice with AST-004 alone significantly reduced the mean lesion volume to 49.63 +/-7.72% (n = 5, p < 0.001) of control (100 +/-7.12%, n = 7), 24 hours post-stroke. In mice treated with FAc, however, AST-004 did not significantly reduce lesion sizes (82.73 +/-16.69%, n = 5, p = 0.369) (**Fig. 5**). Mice treated with vehicle and FAc exhibited an average lesion size of 65.7 +/-13.11% (n = 5), not significantly different from control mice (**Fig. 5**). We conclude from these data that the cerebroprotective efficacy of AST-004 is dependent on astrocyte mitochondrial metabolism.

**Figure 5:**
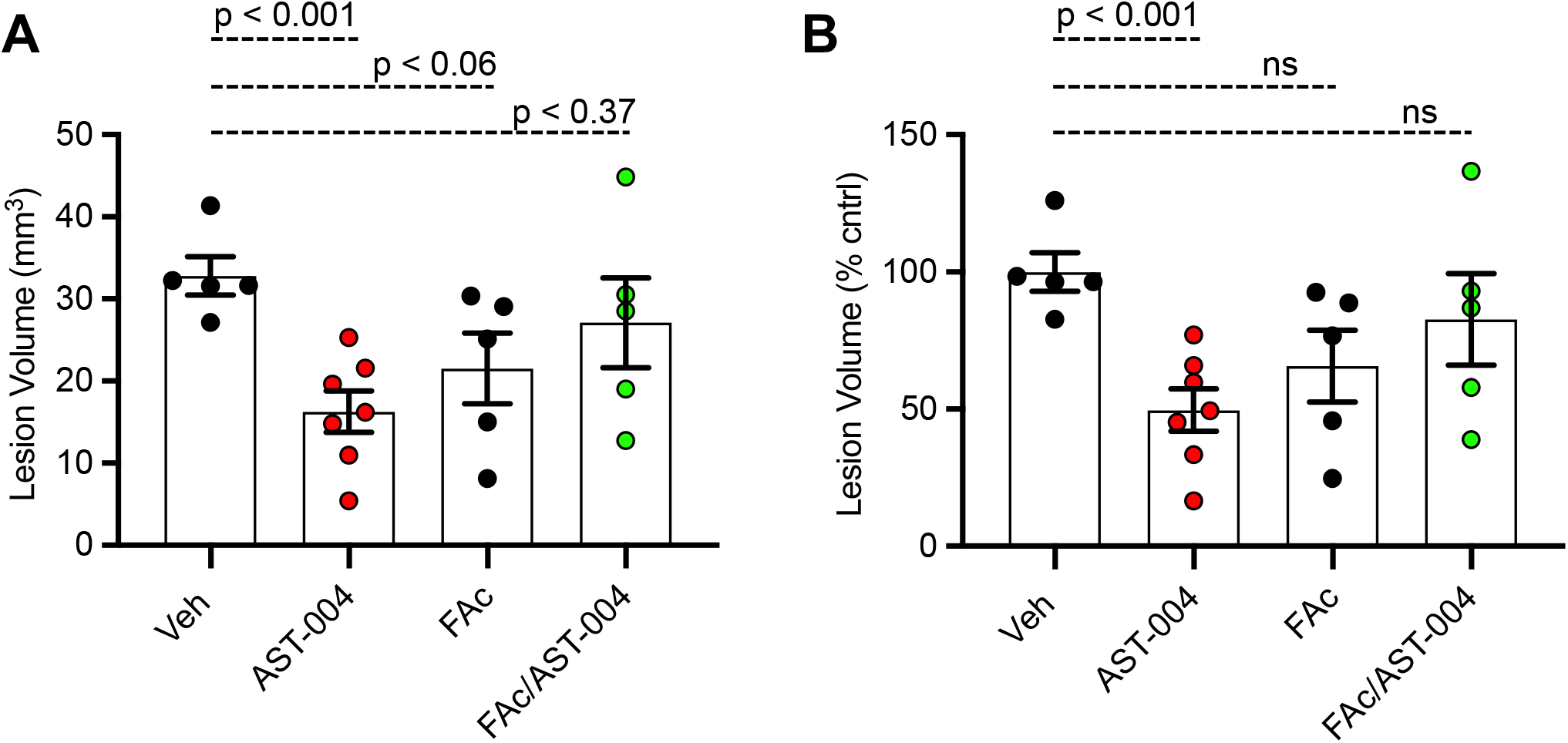
Fluoroacetate, an astrocyte specific mitochondrial toxin, blocks AST-004 neuroprotection. **(A)** Histogram plot of average TTC-stained stroke volumes in mice treated with either vehicle, AST-004, fluoroacetate (FAc) or fluoroacetate and AST-004 (FAc/AST-004). **(B)** Data presented in (A) replotted as a percentage of the average control stroke volume. Data were pooled from 2 experiments using male mice and plotted as mean +/-SEM. ns not significant.

### Photothrombotic stroke increases mRNA levels for *Adora3* and *Gfap* in the areas surrounding the lesion

Numerous reports have indicated A3R expression is upregulated in the context of hypoxia and inflammation^33-35^. To determine if similar changes occur with our model of stroke, we isolated brain tissue surrounding the lesion 24 hours post-stroke for qPCR measurements. We examined mRNA levels for both Adora3 and Gfap, as a marker for inflammation, and compared the relative levels between ipsilateral and contralateral samples. We found a 50% increase in ipsilateral levels of *Adora3* mRNA (1.50 +/-0.05, p < 0.0001, n = 8) compared to contralateral brain tissue in untreated mice (1.00 +/-0.06, n=8) (**Fig. 6 A**). We also found ipsilateral levels of *Gfap* mRNA (4.88 +/-0.24, p < 0.0001, n = 8) increased nearly 5-fold relative to contralateral tissue (1.00+/-0.07, n=8), consistent with reactive astrogliosis occurring in non-treated injured mice (**Fig. 6 B**).

**Figure 6:**
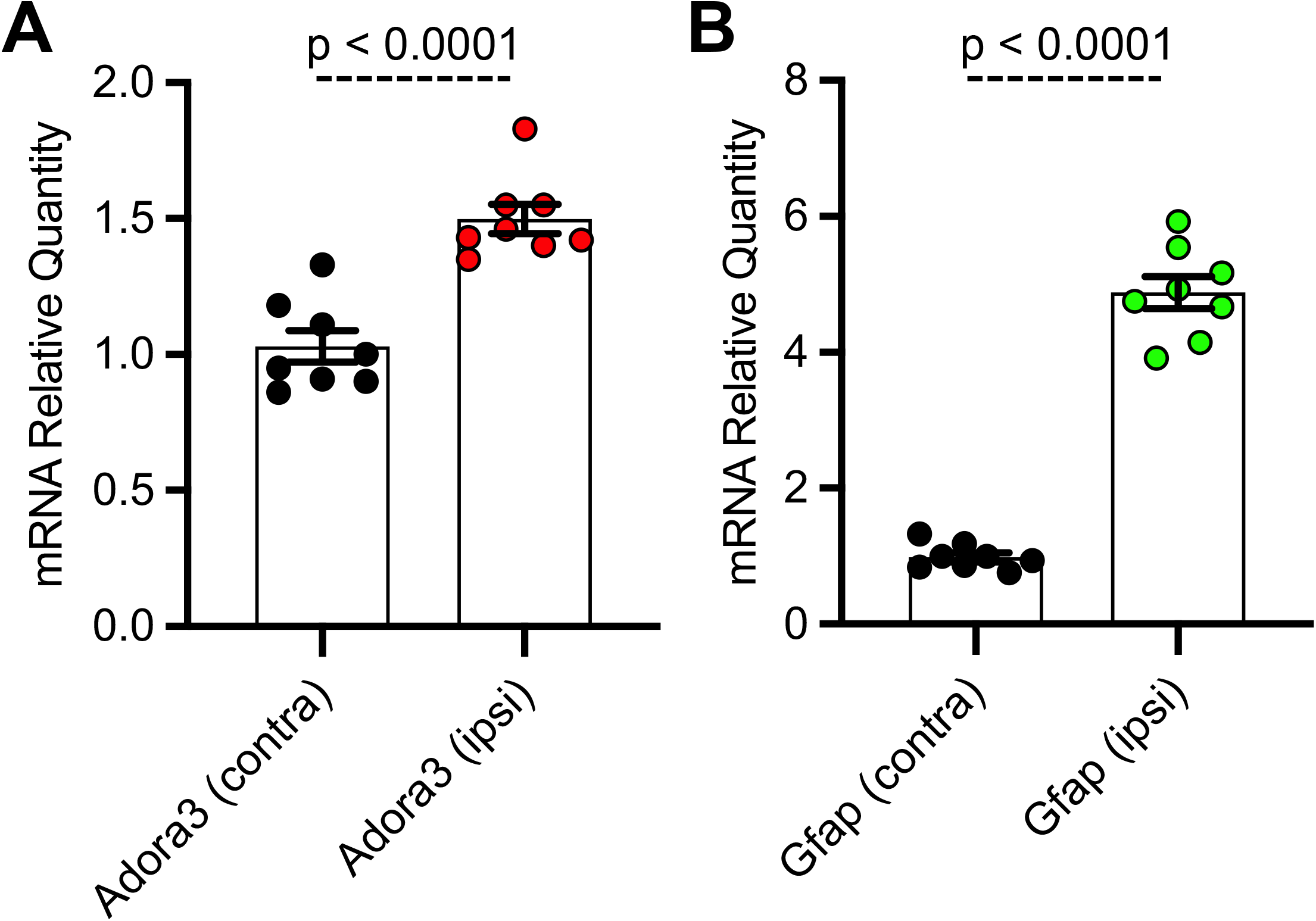
Ipsilateral mRNA levels of *Adora3* and *Gfap* are significantly increased 24 hours post-stroke. **(A)** Relative quantity of *Adora3* mRNA in tissue surrounding photothrombotic lesion site. **(B)** Relative quantity of *Gfap* mRNA in tissue surrounding photothrombotic lesion site. Mice were sacrificed 24 hours post-stroke, their brains removed, mRNA extracted and prepped for qRT-PCR as described in text.

### AST-004 is cerebroprotective in rats after transient middle cerebral artery occlusion (tMCAO)

Given the efficacy of AST-004 in our photothrombotic mouse model of stroke, we also tested whether our A3R agonist was an effective treatment in a transient rat model of stroke. For these experiments, we induced stroke using middle cerebral artery occlusion (MCAO) for 90 minutes followed by reperfusion. AST-004 was intravenously injected at the start of reperfusion followed by constant rate infusions to maintain targeted concentrations through the course of the evaluation period. Three doses of AST-004, a full log difference from each other, were tested. Twenty-four (24) hours post-stroke, we evaluated the functional activities of rats using the modified neurological rating scale (mNRS)^30^. For this assay, no neurologic deficit is scored 0, a failure to extend the left forepaw is scored 1, circling to the left is scored 2, falling to the left is scored 3. Rats that do not walk spontaneously are scored 4 and rats that die are scored 5. For untreated rats, 83% scored between 3 and 5 with a mean mNRS of 3.67 +/-0.36 (n = 12) (**Fig. 7 A**). For rats treated with the mid-dose of AST-004, only 25% scored between 3 and 5, resulting a significantly lower mean mNRS score of 2.25 +/-0.30 (n = 12, p<0.006). Average mNRS scores for the low-dose and high-dose of AST-004 were 3.42 +/-0.38 (n = 12) and 3.00 +/-0.48 (n = 12), respectively. Neither of these mNRS scores were significantly different from untreated rats. Immediately after mNRS scoring, brains were harvested, coronally sliced into 2 mm sections, then stained with TTC as described for mouse brains. The infarct volume for untreated rats, 24 hours post-stroke, was 254.9 +/-33.58 mm^3^ (n = 9) (**Fig. 7 B**). We found AST-004 treatments at the mid-dose (M) significantly (p < 0.005) reduced the mean stroke size to 125.3 +/-36.75 mm^3^ (n = 12). The average lesion size in rats treated with the low-dose (L) of AST-004 was 263.8 +/-44.53 mm^3^ (n = 12) and for rats treated with the high-dose of AST-004, the average lesion size was 213.8 +/-43.90 mm^3^ (n = 12). Neither of these averages were significantly different from untreated rats. Essentially identical findings were observed when infarct volume measurements were replotted as a percentage of the uninjured hemisphere size (**Fig. 7 C**).

**Figure 7:**
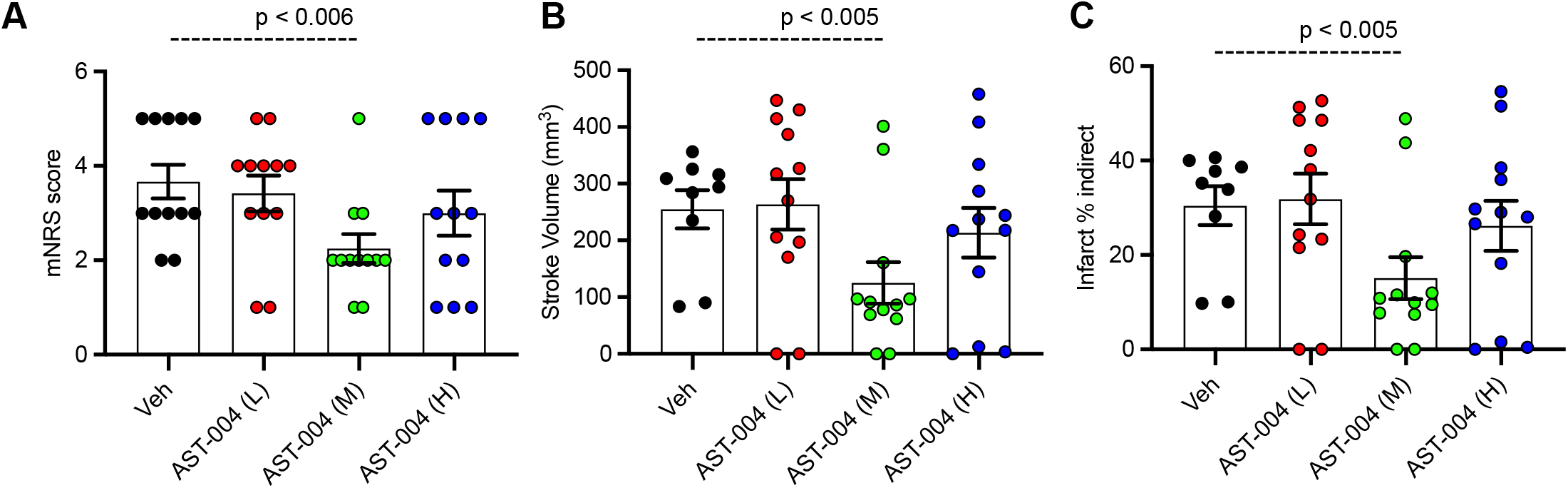
Dose-response of AST-004 treatments for MCAO stroke infarctions in rats. **(A)** Histogram of mNRS scores of animals at each dose of AST-004 tested. **(B)** Average TTC-stained stroke volumes in rats treated with either vehicle or AST-004 at the concentrations labeled: 0.04 mg/kg (L), 0.4 mg/kg (M) and 3 mg/kg (H). **(C)** Indirect measurement of lesion volume through comparative measurements of contralateral hemisphere at doses of AST-004 labeled. Data were collected from male mice and plotted as mean +/-SEM.

## Discussion

Our group previously demonstrated significant cerebroprotection after brain injuries using P2Y_1_R agonists MRS2365 and 2-MeSADP^10,11,13^. Subsequent work showed these phosphorylated nucleotides were rapidly metabolized *in vivo* and that the active cerebroprotective compounds were likely the metabolites of MRS2365 and 2MeSADP, AST-004 and 2-methylthioadenosine, respectively^12^. We confirmed AST-004 treatments were cerebroprotective after TBI in mice^26^ and tMCAO in non-human primates^14^. Binding studies showed AST-004 was primarily an A3R agonist with some affinity for A1Rs^12^.

Here, we validated the cerebroprotective efficacy of AST-004 following photothrombotic stroke in mice, a permanent model of ischemia, as well as following tMCAO in rats, a transient model of ischemia. Efficacy in both permanent and transient ischemia could be of significant clinical importance patients that achieve blood clot removal from either thrombectomy and/or thrombolysis, the current standard of care, and for patients who do not. We also found no sex-dependent effects of this AST-004. This is an important finding considering the often-reported differential effects drugs can have on men and women in clinical settings. Women are known to have a higher incidence of stroke, larger infarcts and worse outcomes, which may be due, in part, to the fact women live longer than men^36^. However, when similarly treated, for example with tissue plasminogen activator (tPA), the difference in outcome measure is significantly reduced^37,38^. The higher incidence of strokes in women may also be affected by the loss of estrogen after menopause^39,40^. Lesion sizes in ovariectomized rodent models are significantly larger, consistent with cerebroprotective effects of estrogens^41^. In our experiments, the average lesion size in untreated female mice trended (p < 0.078) smaller than untreated male mice. Hence, it is possible our female mice benefited from endogenous actions of estrogen. Importantly, when normalized by the mean lesion size for vehicle treated mice, AST-004 significantly decreased lesion size in both females and males.

AST-004 is a lower affinity A1R/A3R agonist that exhibits significant efficacy as shown here and reported previously^14,26^. The cerebroprotective benefits of AST-004 were completely blocked by the A3R antagonist MRS1523, suggesting A1R agonism is not required. Receptor binding models indicate significant efficacy is observed at estimated brain receptor occupancy levels as low as 5% in a non-human primate model of stroke^14^. In light of these results, we tested AST-004 efficacy at doses an order of magnitude lower and higher than previously reported in mice^26^. All AST-004 concentrations were cerebroprotective against stroke, and the lower and higher doses were not significantly different from the mid-dose in mice. In rats, we observed a pronounce U-shaped dose response for both infarct size and the observed neurological deficits. Hormesis, or a “U-Shaped” biphasic dose-response has been observed with many CNS-active agents^42,43^. Importantly, no hormesis was observed in primate stroke efficacy studies, in which a clear AST-004 dose- and concentration-related effect was observed on inhibition of stroke lesion growth^14^.

The observed hormesis in the rat stroke studies, not seen in either mouse or non-human primate studies under our experimental conditions, could be due to the dual affinities of AST-004 for A1Rs and A3Rs. Agonism of both adenosine receptor subtypes can be cerebroprotective in models of ischemic stroke^43-47^. However, A1R agonism has also been associated with cardiovascular effects including bradycardia and hypotension^48,49^. Hypotension has been demonstrated to lead to higher stroke volumes and worse clinical outcomes in ischemic stroke as well as TBI. It is possible that a balance of AST-004 A1R and A3R agonism in rats results in significant efficacy, but at high doses, the peripheral cardiovascular effects of A1R agonism reduce this cerebroprotection. Again, this appears to be a rat-specific phenomenon, since we did not observe any evidence of hormesis or blood pressure effects over a broad dose range in a recent primate stroke efficacy study^14^. More research is required to test the role of A1R agonism at higher doses in our stroke models.

Despite the significantly higher and more specific affinity of MRS5698 and Cl-IB-MECA for the A3R, those compounds had only similar or lower efficacy than AST-004 in the mouse photothrombotic stroke model. This may be due to the pharmacokinetic effects of the significant physicochemical differences between these compounds. AST-004 is a hydrophilic, polar small molecule whereas both MRS5698 and Cl-IB-MECA have substantially higher lipophilicity. These physicochemical differences lead to substantially higher plasma protein and brain tissue binding for MRS5698^44^ and Cl-IB-MECA with correspondingly lower unbound fractions of compound required for distribution and receptor engagement compared to AST-004 (unpublished data). Thus, despite their high affinity for A3R, there are extremely low unbound fractions of these compounds in the brain for A3R agonism. In addition, previous data have suggested that both MRS5698 and Cl-IB-MECA may be excellent substrates for efflux transporters such as P-glycoprotein, potentially further limiting their distribution into the brain^44,45^. AST-004 is not a substrate for P-glycoprotein (unpublished data).

AST-004 cerebroprotection was blocked by the astrocyte specific mitochondrial toxin, fluoroaceteate. Fluoroacetate is preferentially transported into astrocytes by monocarboxylic acid transporter isoforms that are not present in neurons. Its toxic metabolite, fluorocitrate, inhibits the tricarboxylic acid cycle^32^. These data are consistent with previous work showing cerebroprotection with the pro-drug MRS2365 was dependent on astrocyte mitochondrial metabolism^10,11^. In addition, our group demonstrated cerebroprotection by these pro-drugs could be blocked by knocking out the astrocyte specific IP_3_R type2 or by disrupting astrocyte specific mitochondrial function^10,11,46^. Together, these data suggest AST-004 acts by stimulating IP_3_-mediated Ca^2+^ release, leading to enhanced Ca^2+^ sensitive enzyme activity in astrocyte mitochondria. We note that A_3_Rs can effectively couple to either G_q/11_ or G_i/o_ subtypes of G proteins^47,48^.

A3Rs are normally expressed at very low levels in the brain^49-52^. Following photothrombotic stroke, we found a significant increase in brain A3R transcripts, which may aid long-term recovery after brain injury. The availability of additional A3Rs could minimize inherent problems with desensitization from either high levels of endogenous adenosine, which occurs after trauma^53^ or in the presence of an exogenous A3R agonist. Regardless, it is clear mice null for A3Rs exhibit significantly worse outcomes after ischemic injury^16,19,54^. Mechanistically, activation of A3Rs is known to inhibit proinflammatory cytokines^55-57^. Patients with autoimmune inflammatory diseases exhibit high expression of A_3_Rs in peripheral inflammatory cells and in blood mononuclear cells^58,59^. A3Rs are overexpressed in the hypoxic core of tumors and have anti-cancer effects^60,61^. Protein and mRNA levels of A3Rs were also observed to increase after subarachnoid hemorrhage in rats^62^. It is unclear whether our energy-dependent mechanism of cerebroprotection is linked to these anti-inflammatory effects of A3R agonism. Production of reactive oxygen species (ROS) by mitochondria stimulate an inflammatory response^63^. Future studies are needed to test whether ROS production is reduced by AST-004 treatments. Independent, but complimentary mechanisms are also possible. Choi and co-workers reported A3R agonists reduced the lesion volume in rats after MCAO, which significantly decreased recruitment of inflammatory cells to the lesion site^17^.

Overall, we have shown that AST-004 treatments provide significant cerebroprotection in two rodent models of stroke, including treatment of both transient and permanent occlusions. Pharmacological interventions indicate a dependence of this cerebroprotection on A3R agonism and mitochondrial metabolism in astrocytes. Together, these pre-clinal studies confirm the efficacy of AST-004 treatments after brain injuries and encourage the continued development of this new therapeutic in clinical trials.

## References

1. Virani, S.S., Alonso, A., Aparicio, H.J., Benjamin, E.J., Bittencourt, M.S., Callaway, C.W., Carson, A.P., Chamberlain, A.M., Cheng, S., Delling, F.N., Elkind, M.S.V., Evenson, K.R., Ferguson, J.F., Gupta, D.K., Khan, S.S., Kissela, B.M., Knutson, K.L., Lee, C.D., Lewis, T.T., Liu, J., Loop, M.S., Lutsey, P.L., Ma, J., Mackey, J., Martin, S.S., Matchar, D.B., Mussolino, M.E., Navaneethan, S.D., Perak, A.M., Roth, G.A., Samad, Z., Satou, G.M., Schroeder, E.B., Shah, S.H., Shay, C.M., Stokes, A., VanWagner, L.B., Wang, N.Y., Tsao, C.W., American Heart Association Council on, E., Prevention Statistics, C. & Stroke Statistics, S. Heart Disease and Stroke Statistics-2021 Update: A Report From the American Heart Association. Circulation 143, e254–e743 (2021).

2. Collaborators, G.S. Global, regional, and national burden of stroke and its risk factors, 1990-2019: a systematic analysis for the Global Burden of Disease Study 2019. Lancet Neurol. 20, 795–820 (2021).

3. Tissue plasminogen activator for acute ischemic stroke. The National Institute of Neurological Disorders and Stroke rt-PA Stroke Study Group. N. Engl. J. Med. 333, 1581–1587 (1995).

4. Man, S., Zhao, X., Uchino, K., Hussain, M.S., Smith, E.E., Bhatt, D.L., Xian, Y., Schwamm, L.H., Shah, S., Khan, Y. & Fonarow, G.C. Comparison of Acute Ischemic Stroke Care and Outcomes Between Comprehensive Stroke Centers and Primary Stroke Centers in the United States. Circ. Cardiovasc. Qual. Outcomes 11, e004512 (2018).

5. Powers, W.J., Derdeyn, C.P., Biller, J., Coffey, C.S., Hoh, B.L., Jauch, E.C., Johnston, K.C., Johnston, S.C., Khalessi, A.A., Kidwell, C.S., Meschia, J.F., Ovbiagele, B., Yavagal, D.R. & American Heart Association Stroke, C. 2015 American Heart Association/American Stroke Association Focused Update of the 2013 Guidelines for the Early Management of Patients With Acute Ischemic Stroke Regarding Endovascular Treatment: A Guideline for Healthcare Professionals From the American Heart Association/American Stroke Association. Stroke 46, 3020–3035 (2015).

6. Meyers, P.M., Schumacher, H.C., Connolly, E.S., Jr., Heyer, E.J., Gray, W.A. & Higashida, R.T. Current status of endovascular stroke treatment. Circulation 123, 2591–2601 (2011).

7. Parpura, V., Fisher, E.S., Lechleiter, J.D., Schousboe, A., Waagepetersen, H.S., Brunet, S., Baltan, S. & Verkhratsky, A. Glutamate and ATP at the Interface Between Signaling and Metabolism in Astroglia: Examples from Pathology. Neurochem. Res. 42, 19–34 (2017).

8. Tymianski, M. Novel approaches to neuroprotection trials in acute ischemic stroke. Stroke 44, 2942–2950 (2013).

9. Grupke, S., Hall, J., Dobbs, M., Bix, G.J. & Fraser, J.F. Understanding history, and not repeating it. Neuroprotection for acute ischemic stroke: from review to preview. Clin. Neurol. Neurosurg. 129, 1–9 (2015).

10. Zheng, W., Watts, L.T., Holstein, D.M., Prajapati, S.I., Keller, C., Grass, E.H., Walter, C.A. & Lechleiter, J.D. Purinergic receptor stimulation reduces cytotoxic edema and brain infarcts in mouse induced by photothrombosis by energizing glial mitochondria. PloS one 5, e14401 (2010).

11. Zheng, W., Talley Watts, L., Holstein, D.M., Wewer, J. & Lechleiter, J.D. P2Y1R-initiated, IP3R-dependent stimulation of astrocyte mitochondrial metabolism reduces and partially reverses ischemic neuronal damage in mouse. J. Cereb. Blood Flow Metab. 33, 600–611 (2013).

12. Liston, T.E., Hinz, S., Muller, C., Holstein, D., Wendling, J., Melton, R.J., Campbell, M., Korinek, W.S., Suresh, R.R., Sethre-Hofstad, D., Gao, Z.G., Tosh, D.K., Jacobson, K.A. & Lechleiter, J.D. Nucleotide P2Y1 Receptor Agonists are In Vitro and In Vivo Prodrugs of A1/A3 Adenosine Receptor Agonists: Implications for Roles of P2Y1 and A1/A3 Adenosine Receptors in Health and Disease. Purinergic Signal in press(2020).

13. Talley Watts, L., Sprague, S., Zheng, W., Garling, R.J., Jimenez, D., Digicaylioglu, M. & Lechleiter, J. Purinergic 2Y(1) receptor stimulation decreases cerebral edema and reactive gliosis in a traumatic brain injury model. J. Neurotrauma 30, 55–66 (2013).

14. Liston, T.E., Hama, A., Boltze, J., Poe, R.B., Natsume, T., Hayashi, I., Takamatsu, H., Korinek, W.S. & Lechleiter, J.D. Adenosine A1R/A3R (Adenosine A1 and A3 Receptor) Agonist AST-004 Reduces Brain Infarction in a Nonhuman Primate Model of Stroke. Stroke 53, 238–248 (2022).

15. Bjorklund, O., Shang, M., Tonazzini, I., Dare, E. & Fredholm, B.B. Adenosine A1 and A3 receptors protect astrocytes from hypoxic damage. Eur. J. Pharmacol. 596, 6–13 (2008).

16. Chen, G.J., Harvey, B.K., Shen, H., Chou, J., Victor, A. & Wang, Y. Activation of adenosine A3 receptors reduces ischemic brain injury in rodents. J. Neurosci. Res. 84, 1848–1855 (2006).

17. Choi, I.Y., Lee, J.C., Ju, C., Hwang, S., Cho, G.S., Lee, H.W., Choi, W.J., Jeong, L.S. & Kim, W.K. A3 adenosine receptor agonist reduces brain ischemic injury and inhibits inflammatory cell migration in rats. Am. J. Pathol. 179, 2042–2052 (2011).

18. Choi, J.W., Yoo, B.K., Ryu, M.K., Choi, M.S., Park, G.H. & Ko, K.H. Adenosine and purine nucleosides prevent the disruption of mitochondrial transmembrane potential by peroxynitrite in rat primary astrocytes. Arch. Pharm. Res. 28, 810–815 (2005).

19. Fedorova, I.M., Jacobson, M.A., Basile, A. & Jacobson, K.A. Behavioral characterization of mice lacking the A3 adenosine receptor: sensitivity to hypoxic neurodegeneration. Cell. Mol. Neurobiol. 23, 431–447 (2003).

20. Rudolphi, K.A. & Schubert, P. Adenosine and brain ischemia. in Adenosine and ademime nucleotides: from molecular biology to integrative physiology (eds. Belardinell, L. & Pelleg, A.) 391–396 (Kluwer Norwell, 1995).

21. Rudolphi, K.A., Schubert, P., Parkinson, F.E. & Fredholm, B.B. Neuroprotective role of adenosine in cerebral ischaemia. Trends Pharmacol. Sci. 13, 439–445 (1992).

22. von Lubitz, D.K., Carter, M.F., Beenhakker, M., Lin, R.C. & Jacobson, K.A. Adenosine: a prototherapeutic concept in neurodegeneration. Ann. N. Y. Acad. Sci. 765, 163–178; discussion 196-167 (1995).

23. Von Lubitz, D.K., Lin, R.C., Boyd, M., Bischofberger, N. & Jacobson, K.A. Chronic administration of adenosine A3 receptor agonist and cerebral ischemia: neuronal and glial effects. Eur. J. Pharmacol. 367, 157–163 (1999).

24. Von Lubitz, D.K., Lin, R.C., Popik, P., Carter, M.F. & Jacobson, K.A. Adenosine A3 receptor stimulation and cerebral ischemia.. Eur. J. Pharmacol 263, 59–67 (1994).

25. Von Lubitz, D.K., Simpson, K.L. & Lin, R.C. Right thing at a wrong time? Adenosine A3 receptors and cerebroprotection in stroke. Ann. N. Y. Acad. Sci. 939, 85–96 (2001).

26. Bozdemir, E., Vigil, F.A., Chun, S.H., Espinoza, L., Bugay, V., Khoury, S.M., Holstein, D.M., Stoja, A., Lozano, D., Tunca, C., Sprague, S.M., Cavazos, J.E., Brenner, R., Liston, T.E., Shapiro, M.S. & Lechleiter, J.D. Neuroprotective Roles of the Adenosine A3 Receptor Agonist AST-004 in Mouse Model of Traumatic Brain Injury. Neurotherapeutics (2021).

27. Farr, S.A., Cuzzocrea, S., Esposito, E., Campolo, M., Niehoff, M.L., Doyle, T.M. & Salvemini, D. Adenosine A3 receptor as a novel therapeutic target to reduce secondary events and improve neurocognitive functions following traumatic brain injury. J. Neuroinflammation 17, 339 (2020).

28. Ravi, R.G., Kim, H.S., Servos, J., Zimmermann, H., Lee, K., Maddileti, S., Boyer, J.L., Harden, T.K. & Jacobson, K.A. Adenine nucleotide analogues locked in a Northern methanocarba conformation: enhanced stability and potency as P2Y(1) receptor agonists. J. Med. Chem. 45, 2090–2100 (2002).

29. Stubblefield, J.J., Gao, P., Kilaru, G., Mukadam, B., Terrien, J. & Green, C.B. Temporal Control of Metabolic Amplitude by Nocturnin. Cell Rep. 22, 1225–1235 (2018).

30. Longa, E.Z., Weinstein, P.R., Carlson, S. & Cummins, R. Reversible middle cerebral artery occlusion without craniectomy in rats. Stroke 20, 84–91 (1989).

31. Dunwiddie, T.V. & Masino, S.A. The role and regulation of adenosine in the central nervous system. Annu. Rev. Neurosci. 24, 31–55 (2001).

32. Fonnum, F., Johnsen, A. & Hassel, B. Use of fluorocitrate and fluoroacetate in the study of brain metabolism. Glia 21, 106–113 (1997).

33. Godini, R., Fallahi, H. & Ebrahimie, E. Network analysis of inflammatory responses to sepsis by neutrophils and peripheral blood mononuclear cells. PLoS One 13, e0201674 (2018).

34. Morschl, E., Molina, J.G., Volmer, J.B., Mohsenin, A., Pero, R.S., Hong, J.S., Kheradmand, F., Lee, J.J. & Blackburn, M.R. A3 adenosine receptor signaling influences pulmonary inflammation and fibrosis. Am. J. Respir. Cell Mol. Biol. 39, 697–705 (2008).

35. Torres, A., Erices, J.I., Sanchez, F., Ehrenfeld, P., Turchi, L., Virolle, T., Uribe, D., Niechi, I., Spichiger, C., Rocha, J.D., Ramirez, M., Salazar-Onfray, F., San Martin, R. & Quezada, C. Extracellular adenosine promotes cell migration/invasion of Glioblastoma Stem-like Cells through A3 Adenosine Receptor activation under hypoxia. Cancer Lett. 446, 112–122 (2019).

36. Girijala, R.L., Sohrabji, F. & Bush, R.L. Sex differences in stroke: Review of current knowledge and evidence. Vasc. Med. 22, 135–145 (2017).

37. Berglund, A., Schenck-Gustafsson, K. & von Euler, M. Sex differences in the presentation of stroke. Maturitas 99, 47–50 (2017).

38. Ahnstedt, H., McCullough, L.D. & Cipolla, M.J. The Importance of Considering Sex Differences in Translational Stroke Research. Transl Stroke Res 7, 261–273 (2016).

39. Pabbidi, M.R., Kuppusamy, M., Didion, S.P., Sanapureddy, P., Reed, J.T. & Sontakke, S.P. Sex differences in the vascular function and related mechanisms: role of 17beta-estradiol. Am. J. Physiol. Heart Circ. Physiol. 315, H1499–H1518 (2018).

40. Franconi, F., Campesi, I., Occhioni, S. & Tonolo, G. Sex-gender differences in diabetes vascular complications and treatment. Endocr. Metab. Immune Disord. Drug Targets 12, 179–196 (2012).

41. Suzuki, S., Brown, C.M. & Wise, P.M. Neuroprotective effects of estrogens following ischemic stroke. Front. Neuroendocrinol. 30, 201–211 (2009).

42. Calabrese, E.J. Dose-response features of neuroprotective agents: an integrative summary. Crit Rev Toxicol 38, 253–348 (2008).

43. Hammarberg, C., Schulte, G. & Fredholm, B.B. Evidence for functional adenosine A3 receptors in microglia cells. Journal of neurochemistry 86, 1051–1054 (2003).

44. Tosh, D.K., Padia, J., Salvemini, D. & Jacobson, K.A. Efficient, large-scale synthesis and preclinical studies of MRS5698, a highly selective A3 adenosine receptor agonist that protects against chronic neuropathic pain. Purinergic Signal 11, 371–387 (2015).

45. Abel, B., Tosh, D.K., Durell, S.R., Murakami, M., Vahedi, S., Jacobson, K.A. & Ambudkar, S.V. Evidence for the Interaction of A3 Adenosine Receptor Agonists at the Drug-Binding Site(s) of Human P-glycoprotein (ABCB1). Mol. Pharmacol. 96, 180–192 (2019).

46. Zheng, W., Talley Watts, L., Sayre, N.L., Holstein, D. & Lechleiter, J.D. Targeting Astrocyte Mitochondrial ATP production as a Strategy to Treat Brain Injuries. in Glial Biology in Medicine (ed. Sontheimer, H.) (Birmingham, AB, 2012).

47. Abbracchio, M.P., Brambilla, R., Ceruti, S., Kim, H.O., von Lubitz, D.K., Jacobson, K.A. & Cattabeni, F. G protein-dependent activation of phospholipase C by adenosine A3 receptors in rat brain. Mol. Pharmacol. 48, 1038–1045 (1995).

48. Gilman, A.G. G proteins: transducers of receptor-generated signals. Annu. Rev. Biochem. 56, 615–649 (1987).

49. Gessi, S., Merighi, S., Varani, K., Leung, E., Mac Lennan, S. & Borea, P.A. The A3 adenosine receptor: an enigmatic player in cell biology. Pharmacol. Ther. 117, 123–140 (2008).

50. Jacobson, K.A., Nikodijevic, O., Shi, D., Gallo-Rodriguez, C., Olah, M.E., Stiles, G.L. & Daly, J.W. A role for central A3-adenosine receptors. Mediation of behavioral depressant effects. FEBS Lett. 336, 57–60 (1993).

51. Ji, X.d., Lubitz, D.K.J.E., Olah, M.E., Stiles, G.L. & Jacobson, K.A. Species differences in ligand affinity at central AS adenosine receptors. Drug Development Research 33(1994).

52. Lopes, L.V., Rebola, N., Pinheiro, P.C., Richardson, P.J., Oliveira, C.R. & Cunha, R.A. Adenosine A3 receptors are located in neurons of the rat hippocampus. Neuroreport 14, 1645–1648 (2003).

53. Bell, M.J., Kochanek, P.M., Carcillo, J.A., Mi, Z., Schiding, J.K., Wisniewski, S.R., Clark, R.S., Dixon, C.E., Marion, D.W. & Jackson, E. Interstitial adenosine, inosine, and hypoxanthine are increased after experimental traumatic brain injury in the rat. J Neurotrauma 15, 163–170 (1998).

54. Cheng, H., Lederer, W.J. & Cannell, M.B. Calcium sparks: elementary events underlying excitation-contraction coupling in heart muscle. Science 262, 740–744 (1993).

55. Wahlman, C., Doyle, T.M., Little, J.W., Luongo, L., Janes, K., Chen, Z., Esposito, E., Tosh, D.K., Cuzzocrea, S., Jacobson, K.A. & Salvemini, D. Chemotherapy-induced pain is promoted by enhanced spinal adenosine kinase levels through astrocyte-dependent mechanisms. Pain 159, 1025–1034 (2018).

56. Tosh, D.K., Finley, A., Paoletta, S., Moss, S.M., Gao, Z.G., Gizewski, E.T., Auchampach, J.A., Salvemini, D. & Jacobson, K.A. In vivo phenotypic screening for treating chronic neuropathic pain: modification of C2-arylethynyl group of conformationally constrained A3 adenosine receptor agonists. J. Med. Chem. 57, 9901–9914 (2014).

57. Jacobson, K.A., Giancotti, L.A., Lauro, F., Mufti, F. & Salvemini, D. Treatment of chronic neuropathic pain: purine receptor modulation. Pain 161, 1425–1441 (2020).

58. Bar-Yehuda, S., Silverman, M.H., Kerns, W.D., Ochaion, A., Cohen, S. & Fishman, P. The anti-inflammatory effect of A3 adenosine receptor agonists: a novel targeted therapy for rheumatoid arthritis. Expert Opin Investig Drugs 16, 1601–1613 (2007).

59. Ochaion, A., Bar-Yehuda, S., Cohen, S., Barer, F., Patoka, R., Amital, H., Reitblat, T., Reitblat, A., Ophir, J., Konfino, I., Chowers, Y., Ben-Horin, S. & Fishman, P. The anti-inflammatory target A(3) adenosine receptor is over-expressed in rheumatoid arthritis, psoriasis and Crohn’s disease. Cell. Immunol. 258, 115–122 (2009).

60. Fishman, P., Bar-Yehuda, S., Madi, L. & Cohn, I. A3 adenosine receptor as a target for cancer therapy. Anticancer Drugs 13, 437–443 (2002).

61. Madi, L., Ochaion, A., Rath-Wolfson, L., Bar-Yehuda, S., Erlanger, A., Ohana, G., Harish, A., Merimski, O., Barer, F. & Fishman, P. The A3 adenosine receptor is highly expressed in tumor versus normal cells: potential target for tumor growth inhibition. Clin. Cancer Res. 10, 4472–4479 (2004).

62. Li, P., Li, X., Deng, P., Wang, D., Bai, X., Li, Y., Luo, C., Belguise, K., Wang, X., Wei, X., Xia, Z. & Yi, B. Activation of adenosine A3 receptor reduces early brain injury by alleviating neuroinflammation after subarachnoid hemorrhage in elderly rats. Aging (Albany N. Y.) 13, 694–713 (2020).

63. Fishman, P., Bar-Yehuda, S., Liang, B.T. & Jacobson, K.A. Pharmacological and therapeutic effects of A3 adenosine receptor agonists. Drug Discov Today 17, 359–366 (2012).

